# Transcriptome analysis of the Nematode *Caenorhabditis elegans* in acidic stress environments

**DOI:** 10.1101/2020.03.06.980102

**Authors:** Yanyi Cong, Hanwen Yang, Pengchi Zhang, Yusu Xie, Xuwen Cao, Liusuo Zhang

## Abstract

Ocean acidification and acid rain, caused by modern industrial fossil fuels burning, lead to decrease of living environmental pH, which results in a series of negative effects on many organisms. However, the underlying mechanisms of animals’response to acidic pH stress are largely unknown. In this study, we used the nematode *Caenorhabditis elegans* as an animal model to explore the regulatory mechanisms of organisms’response to pH decline. Two major stress-responsive pathways were found through transcriptome analysis in acidic stress environments. Firstly, when the pH dropped from 6.33 to 4.33, the worms responded to the pH stress by up-regulation of the *col*, *nas* and *dpy* genes, which are required for cuticle synthesis and structure integrity. Secondly, when the pH continued to decrease from 4.33, the metabolism of xenobiotics by cytochrome P450 pathway genes (*cyp, gst, ugt,* and ABC transporters) played a major role in protecting the nematodes from the toxic substances probably produced by the more acidic environment. At the same time, cuticle synthesis slowed down might due to its insufficient protective ability. Moreover, the systematic regulation pattern we found in nematodes, might also be applied to other invertebrate and vertebrate animals to survive in the changing pH environments. Thus, our data might lay the foundation to identify the master gene(s) responding and adaptation to acidic pH stress in further studies, and might also provide new solutions to improve assessment and monitoring of ecological restoration outcomes, or generate novel genotypes via genome editing for restoring in challenging environments especially in the context of acidic stress through global climate change.

## 1 Introduction

As an essential environmental factor, pH affects many life processes such as growth, development, metabolism, as well as immune regulation. Whether on land or in water, living organisms often experience pH fluctuations. In recent years, increasing carbon dioxide emissions contribute to ocean acidification and increase the challenges of living environment pressures faced by marine organisms. Due to the increase in H^+^, the marine chemical balance will be broken down, making a variety of marine organisms and even ecosystems that depend on the stability of the chemical environment face great threats (Cornwall and Eddy, 2015; Doubleday et al., 2018). For example, a decrease in the pH of seawater can seriously affect the survival and development of larvae of Atlantic cod, causing damage to important organs such as gills and liver (Stiasny et al., 2018). The descend in pH has a negative impact on the metamorphosis of Pacific oyster larvae by down-regulating several proteins involved in energy production, metabolism and protein synthesis (Dineshram et al., 2016).

At the same time, an increase in the acidity of the rain will also cause a drop in the pH of the soil. China has become the third largest zone of acid rain pollution in the world after North America and Europe (Rodhe et al., 2002). In 2018, 18.9% of cities received precipitations with annual average pH lower than 5.6 (Ministry of Ecology and Environment of the People’s Republic of China, 2019), which means that around one in five cities in China are enveloped by acid rains. Soil acidification will change the composition and function of soil biological communities, reduce the diversity of soil animal communities (Liu et al., 2014; Wang et al., 2014; Wei et al., 2017) and inhibit the activities of soil animals (Zhang et al., 2015a). Under low soil pH, seedling roots in wheat up-regulate genes involved in developmental processes, immune system processes, multi-organism processes, positive regulation of biological processes and metabolic processes of the biological processes. Meanwhile, it down-regulates genes belong to the molecular function category including transporter activity, antioxidant activity and molecular transducer activity and to the extracellular region of the cellular components category (Hu et al., 2018).

In addition, pH is one of the most frequent ecological factors in aquaculture. Many factors can lead to a decrease in the pH of the aquaculture water environment, such as the respiration of plants and plankton in the water, the decomposition of organic matter, and the discharge of acid industrial pollutants. Especially in the intensive aquaculture system, the change of pH is inevitable due to the increase of breeding density and the limitation of the water circulation. There is more organic matter accumulated in the deep sludge, hence the pH value of the bottom water is generally the lowest (Delgado et al., 2003). When the pH in the water environment deviates from the appropriate range of aquatic animals, it will cause damage to the tissues and organs of the cultured organisms, resulting in the impact on its energy metabolism, immunity, growth and development (Chen and Chen, 2003; Garcia Parra and Baldisserotto, 2007; Yu et al., 2015), causing serious economic losses to aquaculture industries (Jiang et al., 2016).

As is mentioned above, the phenomenon of pH decline in many environmental changes is pervasive and widespread and will have a series of negative effects on living organisms. Therefore, studying the biology of acidic environment stress is significant. However, the molecular mechanisms of animals’ response to acidic pH stress are largely unknown.

*C. elegans* is widely used as a model organism for many reasons. It is the first multicellular organism to have its genome completely sequenced (*C. elegans* Sequencing Consortium, 1998), which facilitates bioinformatics analysis. Besides, its genetic tractability, ease of handling and maintenance also make *C. elegans* become a powerful model system to study development, neurobiology and stress responses (Brenner, 1974; Corsi et al., 2015; Zhang et al. 2015b; Hunt, 2016). Only two studies were performed to study pH response in *C. elegans*. One reported that *C. elegans* showed a wide range of pH tolerance (pH 3.1~ pH 11.9) (Khanna et al., 1997), the other showed the *cah-4b* transcript, encoding a putative carbonic anhydrase, was up-regulated under alkaline pH (Hall et al., 2008). However, the underlying mechanisms of *C. elegans*’ response to acidic pH stress are mostly mysteries.

In this study, we employed RNA-seq (RNA Sequencing) to ask how *C. elegans* responds to the acidic pH stresses. We found that there might be two major strategies *C. elegans* used to deal with pH stress: firstly, the expression level of cuticle structure and integrity related genes were significantly increased when pH declined from 6.33 to 4.33; secondly, to deal with even lower pH stress, *C. elegans* substantially increased their xenobiotics metabolism by cytochrome P450 pathway genes. Thus, cuticle structure reorganization and xenobiotics metabolism reprogramming might be key pathways in response to pH stress in nematodes. Furthermore, the mechanisms found in this study might be applied to other invertebrate and vertebrate animals. Thus, our data may lay a foundation for further exploring the molecular mechanisms of animals’ response and adaptation to acidic pH stress environment in the wild.

## 2 Methods

### 2.1 Strains

We obtained the *C. elegans* wild-type Bristol N2 strain from the Caenorhabditis Genetics Center (CGC). N2 worms were cultured on nematode growth medium (NGM) agar plates seeded with E. coli OP50 and maintained at 20 °C (Brenner, 1974).

### 2.2 Worm synchronization

In order to obtain synchronized nematodes, the gravid hermaphrodites were treated with basic hypochlorite solution at room temperature until each adult worm was digested (Silva, 2005). Eggs were collected and cultured overnight on unseeded nematode growth medium agar NGM plates at 20 °C until incubation for about 12 hours.

### 2.3 pH viability assay

35 synchronized newly hatched L1 stage N2 nematodes were cultured in seeded 3 cm diameter NGM pH plates (0.3g NaCl, 1.7g Agar, 0.25g Peptone, 97.5ml H_2_O, 0.1ml M MgSO_4_, 0.1ml M CaCl_2_, 0.1ml 5 mg/ml cholesterol in ethanol buffered at the appropriate pH by varying hydrochloric acid and sodium hydroxide accordingly, 2.5ml M K_2_HPO_4_ buffer solution was added to maintain a stable pH). The growth and development of worms were observed and the earliest time of 35 worms’ spawning observed on each pH plate was recorded (pH 2.53, pH 2.73, pH 2.93, pH 3.13, pH 3.33, pH 4.33, pH 5.33, pH 6.33 pH 7.33, pH 8.33, pH 9.33, pH 10.33 and pH 11.33, respectively). Three replicates were performed for each pH conditions.

### 2.4 Preparation of RNA-seq libraries and data analysis

Large-scale cultivation of *C. elegans* with NGM plates until they harbored plenty of eggs, after bleaching, the eggs were allowed to hatch into L1 larvae in 9 cm diameter unseeded NGM plates overnight (Silva, 2005). And then wash the L1 larvae with M9 to 15ml tubes. After centrifugation at 1300 × g for 5 min, the supernatants were discarded. Pipet the L1 larvae sediment (around 60000 worms) to each seeded 9 cm diameter pH plate (pH 2.93, pH 3.13, pH 4.33 and pH 6.33). Three replicates were performed for each pH conditions. Collected the larvae after incubating for 3 hours at 20 °C in different pH environments. The worms were washed with M9 three times to remove the bulk of the residual bacteria. Then the samples were transferred to 1.5ml tubes and the excess supernatants were removed via centrifugation (2000 × g, 5 min). The samples were frozen immediately in liquid nitrogen for 5 min and stored at −80 °C.

After the sample was rapidly ground with liquid nitrogen, RNA was extracted according to the Trizol method. RNA integrity was analyzed by 1% agarose gels electrophoresis and Bioanalyzer 2100 system; RNA purity was checked using the NanoPhotometer® spectrophotometer. Accurate quantification of RNA concentration was measured by the Qubit 2.0 Fluorometer. Sequencing libraries were generated using NEBNext® UltraTM RNA Library Prep Kit for Illumina® (NEB, USA) following manufacturer’s recommendations. qRT-PCR accurately quantifies the effective concentration of the library to ensure library quality. The library were sequenced on an Illumina Hiseq platform and 150 bp paired-end reads were generated. Reference genome and gene model annotation files were downloaded from Wormbase (WS269). Index of the reference genome was constructed using Hisat2 v2.0.5 and paired-end clean reads were aligned with the reference genome using Hisat2 v2.0.5 (Kim et al., 2015; Kim et al., 2019; Pertea et al., 2016). FeatureCounts v1.5.0-p3 was used to count the reads numbers mapped to each gene (Liao et al., 2013). And then FPKM of each gene was calculated based on the length of the gene and reads count mapped to this gene. Differential expression analysis of two groups was performed using the R package DESeq2 (1.16.1) based on the negative binomial distribution (Love et al., 2014). The Benjamini and Hochberg’s approach was used to adjust the resulting P-values to control the false discovery rate. Corrected P-value < 0.05 and Fold Change > 2 were set as the threshold for significantly differential expression. GO enrichment analysis and KEGG pathway enrichment analysis of differentially expressed genes were achieved by clusterProfiler R package with adjusted P-value < 0.05 (Yu et al., 2012).

### 2.5 Quantitative real-time PCR validation

Some of the key genes of our interest were selected for qPCR validation (*col-91, col-146, col-39, nas-38, nas-1, dpy-1, dpy-5, cyp13A5, cyp13A12, gst-31, gst-37, ugt-56, ugt-66, haf-1, mrp-2, nhr-11, nhr-237, hsp-16.1, hsp-70*). Total RNA was obtained by the same method applied in the sample preparation of RNA-seq libraries. Three RNA samples were collected for each pH conditions (pH 2.93, pH 3.13, pH 4.33 and pH 6.33, three biological replicates respectively). The ReverTra Ace qPCR RT Master Mix with gDNA Remover Kit (TOYOBO) was used for reverse transcription. The synthesized cDNA was diluted 1:40. 4 μl of the diluted cDNA was used to carry out qPCR with SYBR® Green Realtime PCR Master mix on a QuantStudio Real-Time PCR System following the manufacturer’s protocol. Primer sequences are provided in Supplementary File 1. Genes tested from each biological replicate had three technical repeats. Each gene of interest was normalized to an internal control gene (*tba-1*) and expressed as a fold change of treatment groups (pH 4.33, pH 3.13, pH 2.93) compared with control group (pH 6.33) respectively. Statistical analysis was performed by applying a two-tailed t-test to compare the treatment groups and control group on GraphPad Prism 5.

## 3 Results

### 3.1 Spawning time of *C. elegans* in different pH environments

To ask if pH stress affect *C. elegans* development and growth, we performed phenotypic experiments in different pH environments. The spawning time of *C. elegans* is an easy-to-observe indicator, so it is feasible to judge the growth rate of *C. elegans* by its spawning time. Our results showed that *C. elegans* is highly resistant to pH stress and it can grow and reproduce in a wide range of pH (3.13-11.33) (Figure 1). The pH of the standard NGM was 6.33, and spawning time was 52 h under this condition; Spawning time delayed to 54 h at pH 4.33 and pH 3.33, which was two hours slower than the control group (pH 6.33); Spawning time was 55 h at pH 3.13, which was 3 hours slower than the control group (Figure 1). Strikingly, pH 2.93 and pH 3.13 only differed by 0.2, but this difference had a huge effect on the growth of N2. We noticed that *C. elegans* cannot lay eggs at pH 2.93, however the nematode can grow and reproduce on the plate of pH 3.13, although with 3 hours delay of eggs laying (Figure 1). The nematode was not able to survive on the pH 2.53 plate, while on the plate of pH 2.73, there are few surviving worms on the 5th day. Furthermore, we found that *C. elegans* had a wide range of adaptation to alkalinity, and the spawning time is about 54h at the range of pH 7.33~11.33 (Figure 1).

**FIGURE 1.**
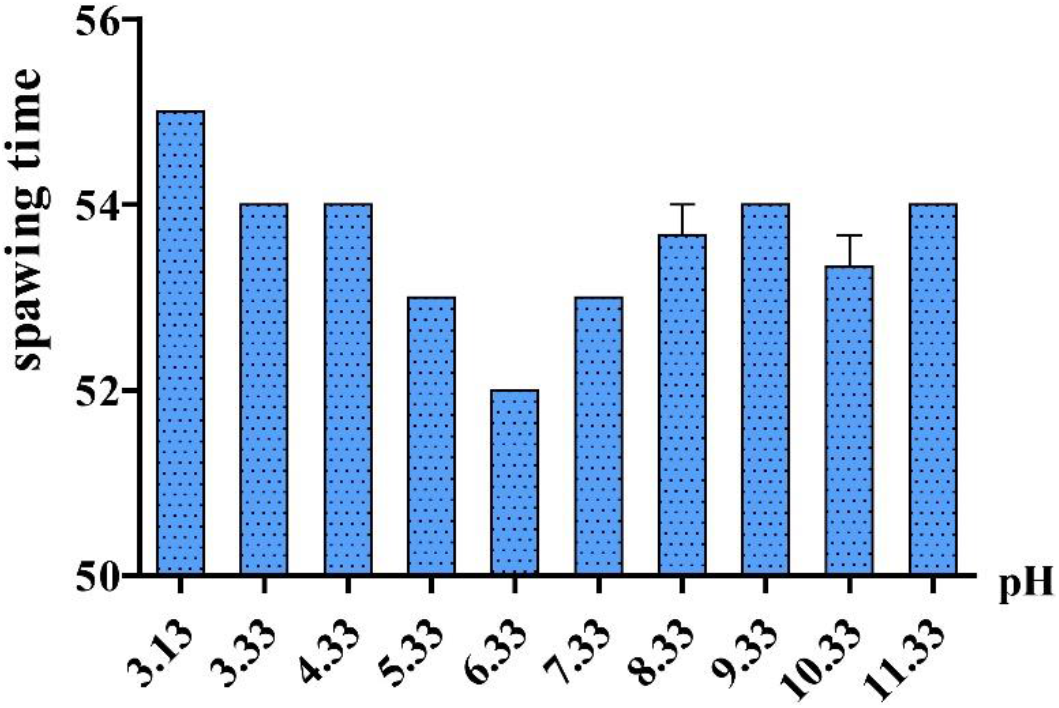
Spawning time in different pH environments. 35 newly hatched L1 were cultured at specific pH medium. The earliest observed spawning time on each plate was recorded. Error bars represent the standard error from replicated experiments (n=3).

### 3.2 Identification of differentially expressed genes in *C. elegans* exposed to acidic environments

In this study, we used high-throughput RNA-seq to identify and quantify the differentially expressed genes (Figure 2). According to the data, we used pH 6.33 as the control group, pH 2.93, pH 3.13, pH 4.33 as the treatment group, FDR < 5% and Fold Change > 2 as the differential gene screening threshold. The number of up-regulated genes and down-regulated genes in each comparison combination is shown in Table1. Details of significantly up-regulated and down-regulated differentially expressed genes in each condition were list in Supplementary File 2 and Supplementary File 3.

**FIGURE 2.**
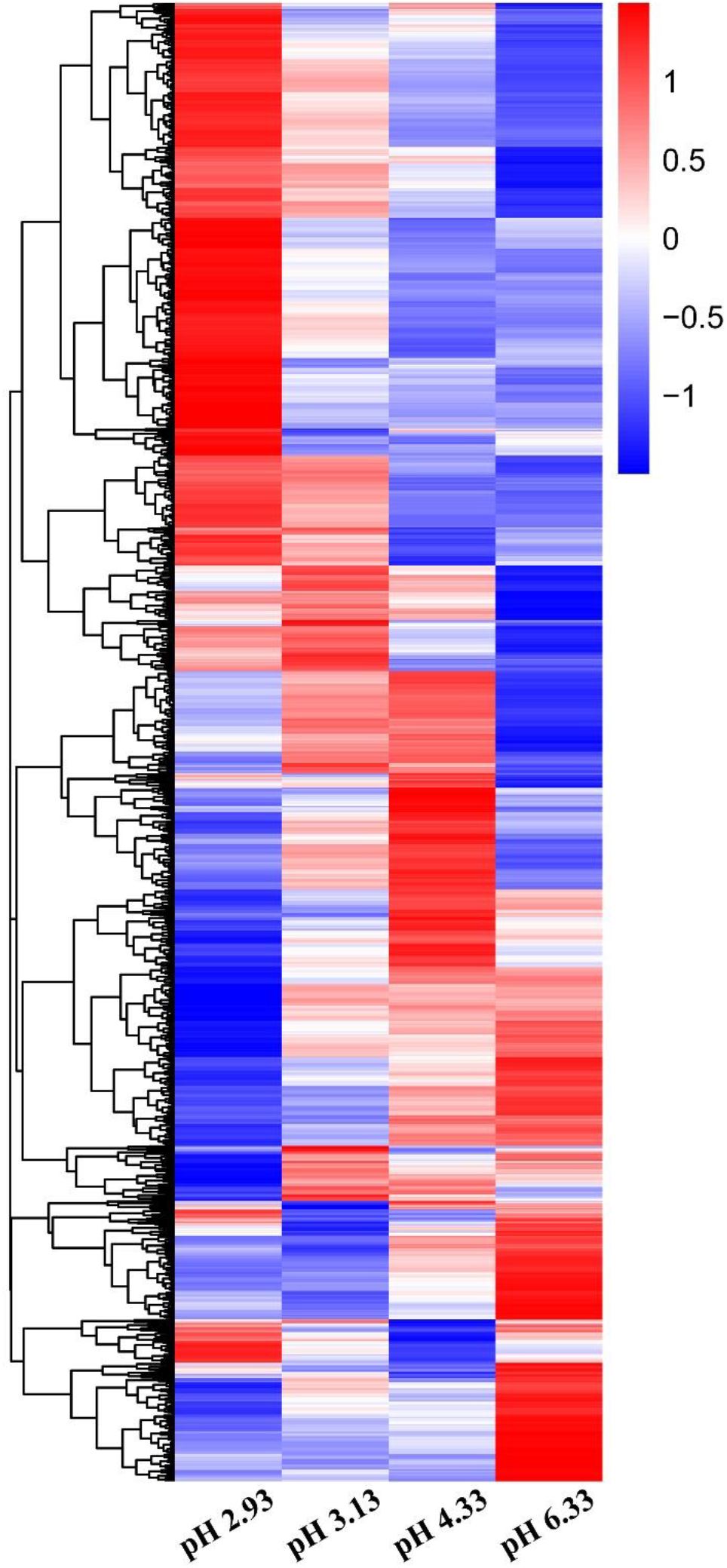
Heatmap of DEGs in *C. elegans* exposed to different acidic pH environments. The scale bar shows the z-score for a differentially expressed gene. Red indicates up-regulation, blue indicates down-regulation.

**TABLE 1.**
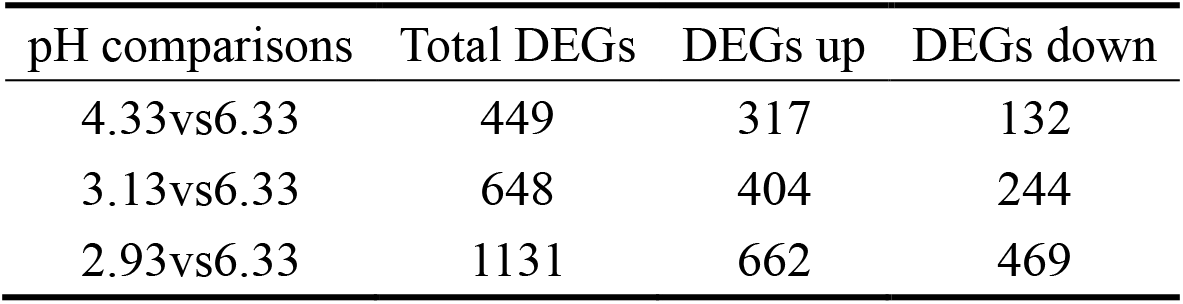
The number of differentially expressed genes.

| pH comparisons | Total DEGs | DEGs up | DEGs down |
| --- | --- | --- | --- |
| 4.33vs6.33 | 449 | 317 | 132 |
| 3.13vs6.33 | 648 | 404 | 244 |
| 2.93vs6.33 | 1131 | 662 | 469 |

DEGs represent ‘differentially expressed genes’. They were identified using DESeq2 package (1.16.1) (FDR < 0.05 and FC > 2)

### 3.3 The expression of epidermal synthesis-related genes increased first and then decreased with the decline of pH

Based on GO analysis, we observed that the expression of genes related to epidermal synthesis varied gradually with the decrease of pH (Supplementary File 4 and Supplementary File 5). *col* and *dpy* encode the collagen genes and are supposed to be structural constituent of cuticle. *nas* encodes an astacin-like metalloprotease and is predicted to have metalloendopeptidase activity (www.wormbase.org). It has been reported that *nas-35* is a gene closely related to epidermal synthesis (Novelli et al., 2004). *nas-1,7,14,23,35* are all expressed in the hypodermis and probably process cuticle components like cuticular collagens (Park et al., 2010). The astacin metalloprotease *nas-36* and *nas-37* play an important role in cuticle ecdysis which result in molting defects when mutated (Stepek et al., 2010). When the pH dropped from 6.33 to 4.33, *col, nas* and *dpy* expression were up-regulated (Figure 3). When the pH continued to decrease from 4.33 to 3.13, the amplitude of the increase in the expression of these three genes decreased. When the pH dropped to 2.93, there was no significant difference compared to the expression levels detected at pH 6.33 (Figure 3). Thus, we propose that cuticle structure and integrity genes play the primary regulatory role to deal with acidic pH stress environments.

**FIGURE 3.**
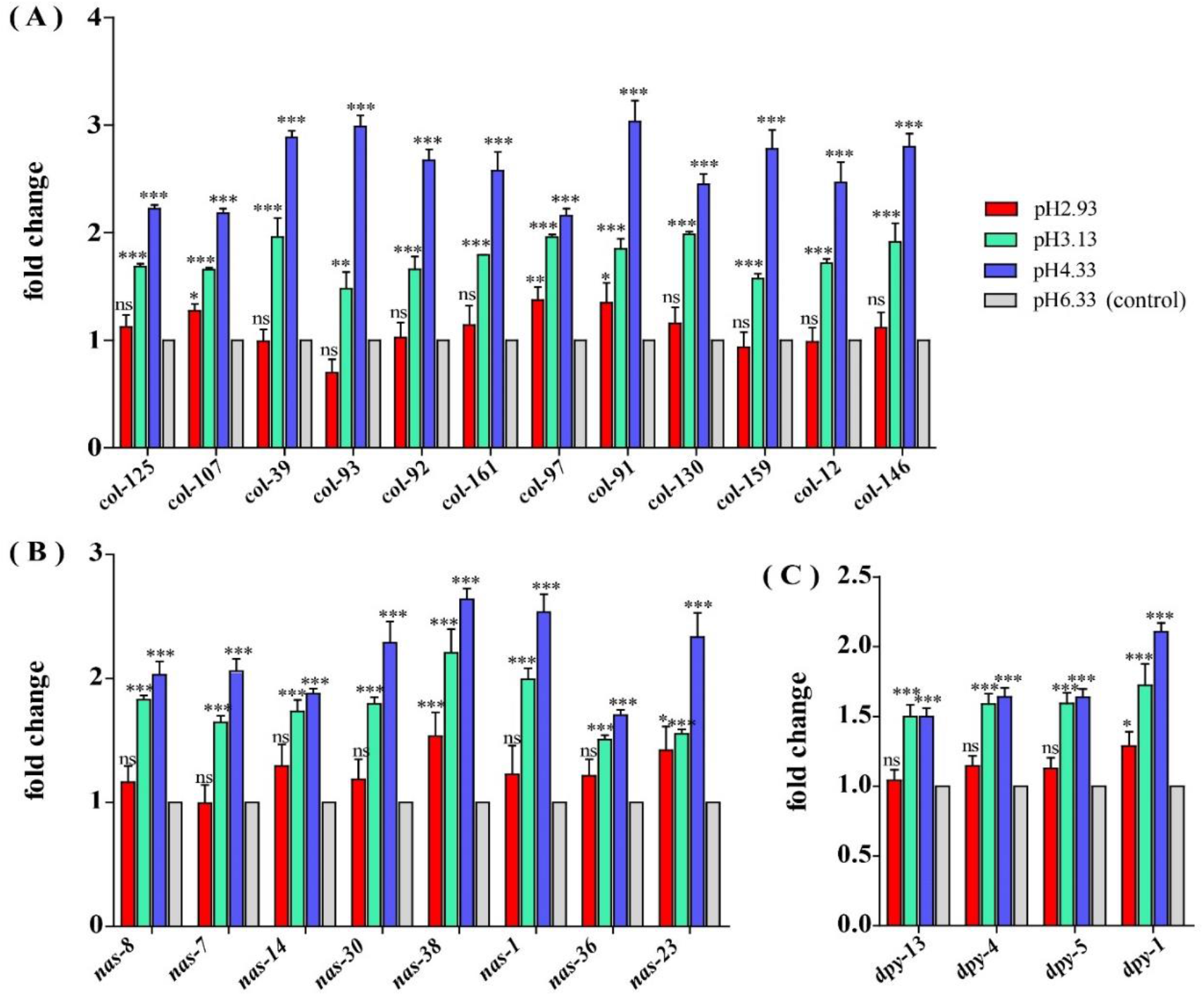
Expression of cuticle synthesis related genes. (A) Transcript level of *col* genes. (B) Transcript level of *nas* genes. (C) Transcript level of *dpy* genes. Fold changes indicate the ratio of the treatment group (pH 2.93, pH 3.13, pH 4.33) to the control group (pH 6.33). The error bars represent standard error of the mean of 3 biological replicates on per condition. \**P* <0.05, \*\**P* <0.01, \*\*\**P* <0.001

### 3.4 Up-regulation genes involved in the metabolism of xenobiotics by cytochrome P450 pathway

Through GO functional enrichment analysis and KEGG pathway analysis, it was found that when the pH decreases from 4.33 to 3.13, the metabolism of xenobiotics by cytochrome P450 pathway was significantly up-regulated (Figure 4, Supplementary File 4 and Supplementary File 5). Cytochrome P450 oxidase is the most important enzyme involved in the first phase of metabolism of xenobiotics, which is an iron porphyrin (erythrocyte heme) protein that catalyzes many types of redox reactions to increases the solubility of a substrate, usually by adding or uncovering a hydrophilic group (such as hydroxyl groups) (Supplementary Figure1)(Barnes, 1960). It can be seen that there was no significant change in the expression level of *cyp* gene when the pH dropped from 6.33 to 4.33 (Figure 5A). However, when the pH dropped to 3.13, the expression of the *cyp* gene was significantly up-regulated (Figure 5A). Especially, we found six *cyp* genes (*cyp-13A8, cyp-33D3, cyp-14A4, cyp-35B2, cyp-13A11, cyp-33C2*), whose expression levels were very low at pH 6.33 and pH 4.33, while those genes were significantly up-regulated when the pH dropped to 3.13 (Figure 5B).

**FIGURE 4.**
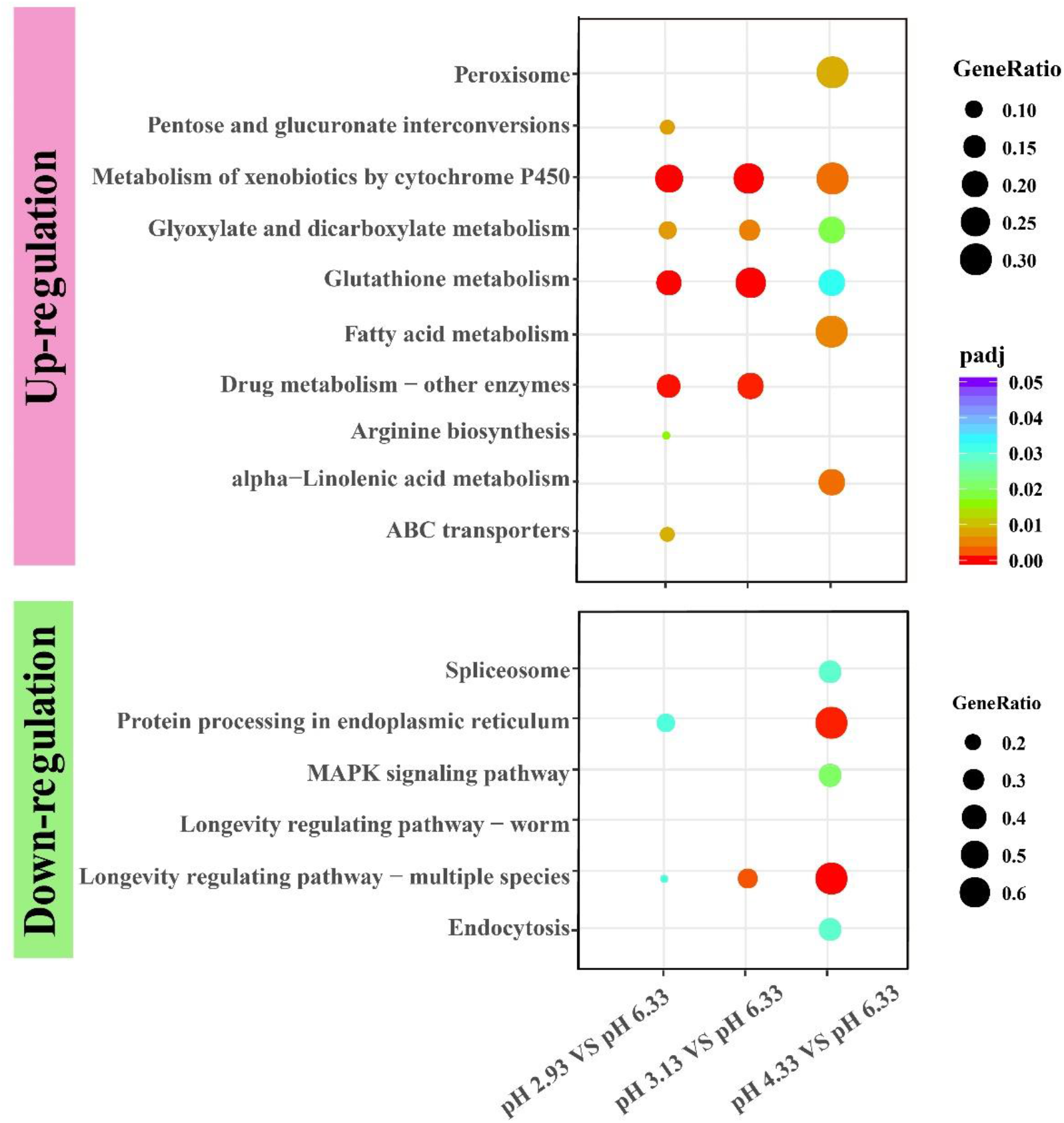
KEGG enrichment of up-regulated and down-regulated genes.

**FIGURE 5.**
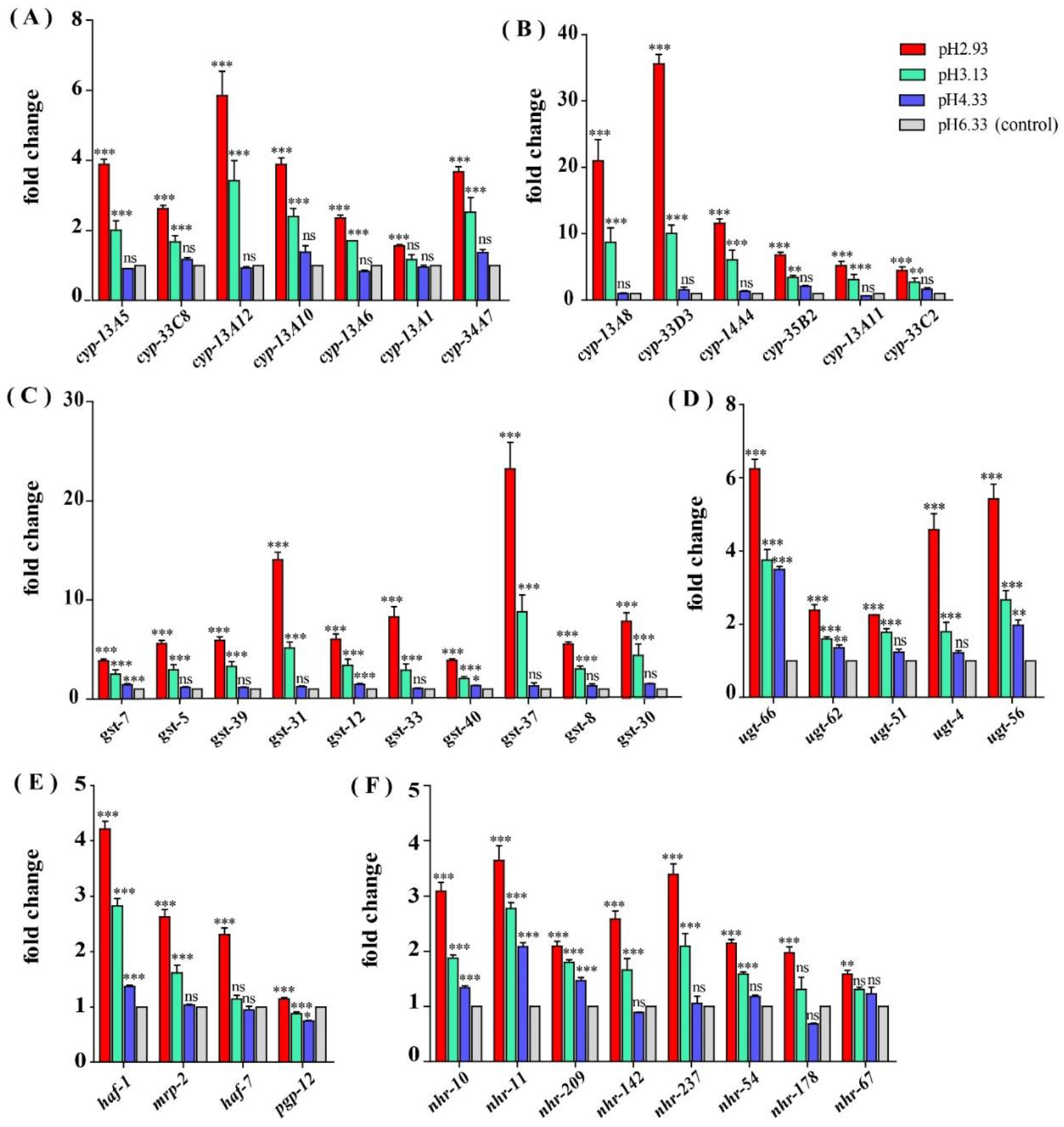
Expression of metabolism of xenobiotics by cytochrome P450 pathway related genes and nuclear hormone receptors genes. (A) Transcript level of *cyp* genes. (B) Transcript level of *cyp* genes whose expression level are very low (hardly expressed) at pH 6.33 and pH 4.33. (C) Transcript level of *gst* genes. (D) Transcript level of *ugt* genes. (E) Transcript level of ABC transporter related genes. (F) Transcript level of nuclear hormone receptor genes. Fold changes indicates the ratio of the treatment group (pH 2.93, pH 3.13, pH 4.33) to the control group (pH 6.33). The error bars represent standard error of the mean of three biological replicates per condition. \**P* <0.05, \*\**P* <0.01, \*\*\**P* <0.001

Similarly, when the pH dropped from 6.33 to 4.33, the expression levels of most *gst* and *ugt* genes weren’t changed significantly (except *gst-7, ugt-56, ugt-62, ugt-66*). However, when the pH dropped to 3.13, the expression levels of 10 *gst* and 5 *ugt* genes were significantly up-regulated (Figure 5C, Figure 5D). UDP-glucuronyltransferase (UGT) and glutathione-S-transferase (GST) are two types of conjugating enzymes during phase II of metabolism of xenobiotics. These two enzymes can catalyze the binding of xenobiotics with polar groups produced in the phase I to glucuronic acid and glutathione respectively, improve the water solubility of the phase I products, which facilitates them to discharge from the body (Supplementary Figure 1) (Croom, 2012).

Besides, the gene expression levels of *haf-1* and *pgp-12* showed an increasing trend with the decreasing pH. When the pH decreased from 6.33 to 4.33, the expression levels of the *mrp-2* and *haf-7* genes didn’t change significantly. However, when the pH dropped to 3.13, the expression level of the *mrp-2* gene was significantly up-regulated. When the pH dropped to 2.93, the expression level of *haf-7* was significantly up-regulated (Figure 5E). *haf-1*, *haf-7*, *mrp-2* and *pgp-12* genes are ABC (ATP-binding cassette) transporters in *C. elegans*. The metabolites that have undergone the first and second phase of reactions are ultimately transported out of the cell by the ABC transporters.

In summary, we found that most xenobiotics detoxification pathway genes increased significantly in severe acidic pH stress environment, including P450 *cyp* genes in phase I, *ugt* and *gst* genes in phase II, and ABC transporters.

### 3.5 Up-regulation of gene expression of nuclear hormone receptors

In addition to the P450 pathway, we found that the gene expression of nuclear hormone receptors also showed a similar trend. It can be seen that the gene expression of *nhr-10*, *nhr-11*, *nhr-209* showed an upward trend when the pH declined (Figure 5F). When the pH dropped from 6.33 to 4.33, the expression levels of *nhr-142*, *nhr-237*, *nhr-54*, and *nhr-178* genes didn’t change significantly. However, when the pH dropped to 3.13, the expression levels of these *nhr* genes were significantly up-regulated (except *nhr-178*).

### 3.6 Down-regulation of heat shock protein gene expression

As a common indicator of cellular stress, heat shock proteins are induced to express under many stimuli such as cold, infiltration, drought, salt, ultraviolet light, oxidative stress and pathogen infection (Swindell et al., 2007). Unexpectedly, our results showed that the expression of heat shock protein genes were significantly down-regulated in acidic pH environment (Figure 6). In this study, these down-regulated heat shock protein genes were enriched into Protein processing in endoplasmic reticulum and Longevity regulated pathway by KEGG pathway analysis (Figure 4). Our results suggested that acidic pH stress might severely affect the capacity of protein processing in *C. elegans*.

**FIGURE 6.**
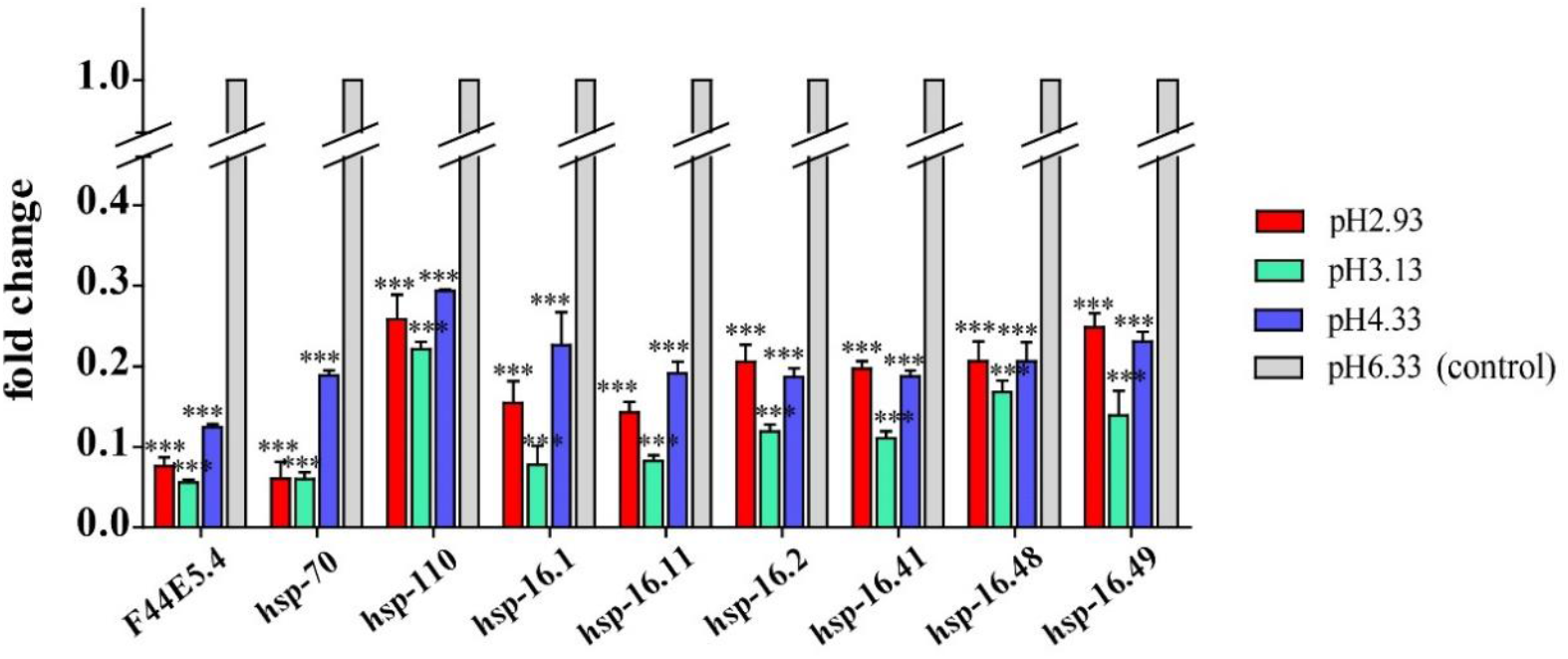
Transcript level of *hsp* genes. Fold changes indicates the ratio of the treatment group (pH 2.93, pH 3.13, pH 4.33) to the control group (pH 6.33). The error bars represent standard error of the mean of three biological replicates per condition. \**P* <0.05, \*\**P* <0.01,\*\*\**P* <0.001

### 3.7 Quantitative real-time PCR validation

Some of the genes of our interest were validated by qPCR. Similar trend were displayed in both the qRT-PCR and RNA-seq analyses. As shown in Figure 7A, expression level of *col*, *nas* and *dpy* genes increased significantly in pH 4.33 and then decreased with the decline of pH. Consistent with our RNA-seq data, expression of *cyp, gst, ugt, haf* and *nhr* genes were increased significantly in pH 3.13 and 2.93 (Figure 7B). In addition, we confirmed that *hsp-70* and *hsp-16.1* genes were down-regulated in all the acidic environments (Figure 7C).

**FIGURE 7.**
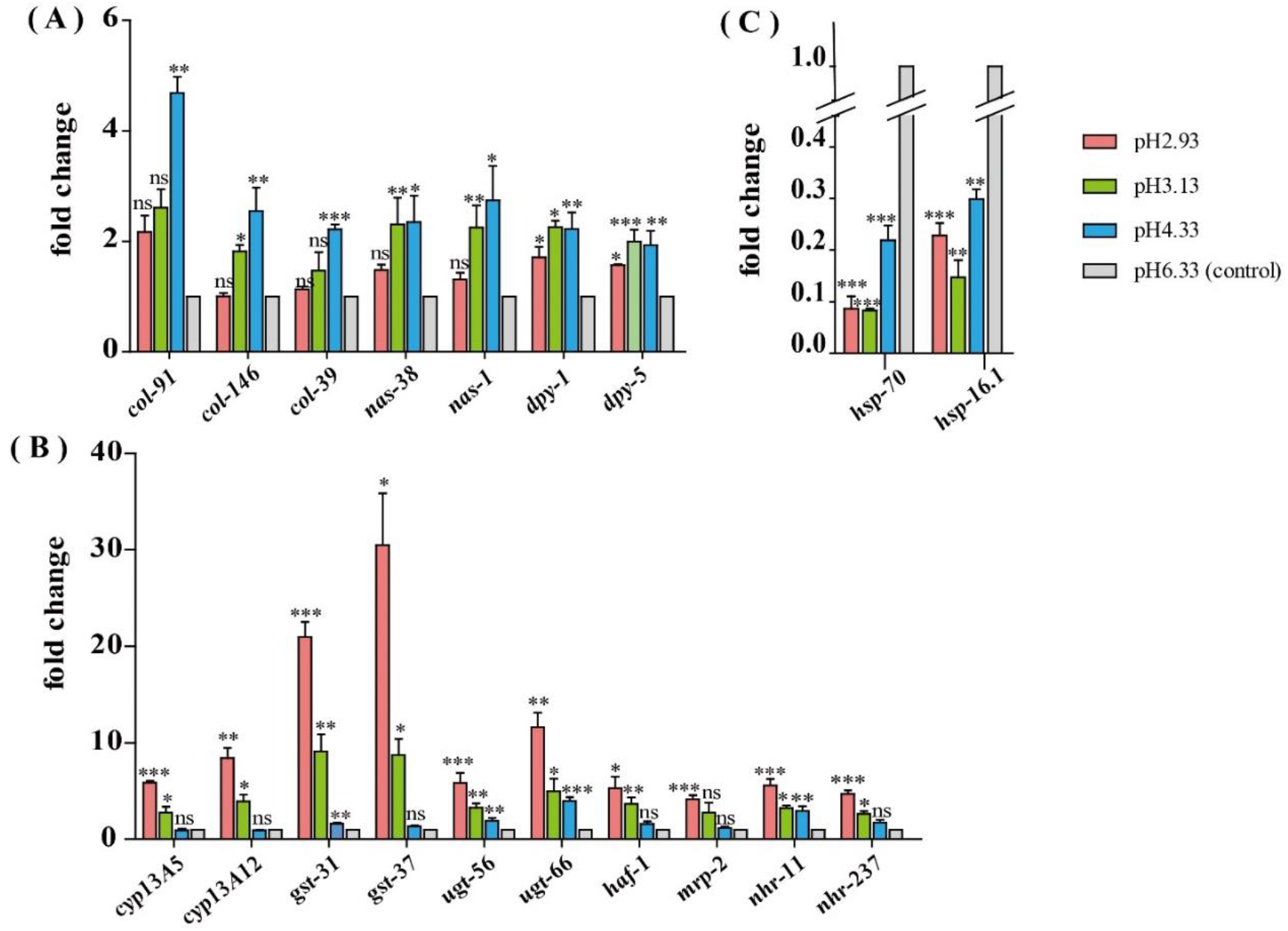
Validation of the RNA-seq results using qRT-PCR. (A) Expression level of cuticle structure and integrity related genes. (B) Expression level of xenobiotics metabolism by cytochrome P450 pathway genes. (C) Expression level of *hsp* genes. Fold change indicates the ratio of the treatment group (pH 2.93, pH 3.13, pH 4.33) to the control group (pH 6.33).The mean fold changes and standard error of the mean of three biological replicates are graphed. \**P* <0.05, \*\**P* <0.01, \*\*\**P* <0.001 (t-test, two-sided)

## 4 Discussion

As an important ecological factor, pH has a direct impact on the life processes of living organisms. Studying the mechanism of pH stress sensation and responding signaling in *C. elegans*, might lay a foundation for understanding how invertebrate animals respond and adapt to environmental pH stresses.

### 4.1 Cuticle remodeling genes are induced by low pH

The nematode cuticle is a highly structured extracellular matrix composed predominantly of cross-linked collagens, additional insoluble proteins termed cuticlins, associated glycoproteins and lipids (Page and Winter, 2003). It forms a hydrostatic skeleton and acts as a primary barrier and first line of defense against many environmental violations. Cuticle collagen synthesis pathway involves numerous co- and post-translational modification, processing, secretion and cross-linking steps that in turn are catalyzed by specific enzymes and chaperones including astacin metalloprotease (Page and Winter, 2003). Nematode astacins are the key to the development of *C. elegans* and have special roles in hatching, molting and cuticle synthesis (Hishida et al., 1996; Page and Winter, 2003; Stepek et al., 2010; Stepek et al., 2015). Loss-of-function mutations in *dpy-31*/*nas-35* lead to typical cuticle synthesis defects (Novelli et al., 2004). The *nas-6*; *nas-7* double mutant grows slowly and has defects in the pharyngeal grinder, which is a cuticular structure important for food processing (Park et al., 2010).

*dpy-2*, *dpy-3*, *dpy-7* and *dpy-10* are necessary for the formation of specific bands of collagen called annular furrows, when these genes are knocked down, annular furrows disappear accordingly and the detoxification, hyperosmotic, and antimicrobial responses will be activated (Dodd et al., 2018). However, the above *dpy* genes didn’t show significantly changes in our study (Supplementary Figure 2). It was reported that *dpy-1*, *dpy-5* and *dpy-13* are required for alae (lateral ridges in the cuticle of the adult *C. elegans*) formation (Dodd et al., 2018; Thein et al., 2003). Surprisingly, significantly expression changes were observed in these alae-related genes in our study, suggesting that the alae might be involved in acidic pH stress response.

Cuticle structure and integrity genes might play primarily regulatory role to deal with acidic pH stress environments. When the pH drops from 6.33 to 4.33, *C. elegans* is likely to fight lower pH stress by enhancing the synthesis and secretion of epidermal collagen, so the expression of *col*, *nas* and *dpy* genes are up-regulated. When the pH continues to decrease from 4.33, the amplitude of the increase in the expression of these genes decreases, which suggests that the damage caused by more acidic environment is too strong to be handled via cuticle physical changes. It has been reported that cuticle structure and integrity, especially annular furrows are damage sensors in certain abiotic and biotic stresses, such as osmotic, xenobiotic and bacterial infection (Dodd et al., 2018).

### 4.2 Metabolism of xenobiotics by cytochrome P450 pathway play a major role in extreme acidic pH stress environments

Our results showed that as the pH decreases, genes involved in the metabolism of xenobiotics by cytochrome P450 pathway were all up-regulated. And the overall trend was that gene expression didn’t produce significant differences when the pH dropped from 6.33 to 4.33. As the pH continued to drop, the expression of genes in this pathway were significantly up-regulated. We propose that the pH decline causes different degrees of acidity pressure on the nematodes, which may lead to the accumulation of toxic substances in the nematode and triggers the up-regulation of multiple genes in the cytochrome P450 pathway. Thus, the cytochrome P450 pathway might be a key metabolic pathway for biological response to severely acidity stress.

When the pH declined from 8.15 to 7.81, the cytochrome p450 like transcript of the great spider crab Hyas araneus was up-regulated by 9.4-fold (Harms et al., 2014). Similarly, Timmins-Schiffman et al. found in the proteomic analysis of the Pacific oysters (Crassostrea gigas) that when the pH fell from 8.0 to 7.3, the expression of Cytochrome P450 1A5 was elevated by 2.7 times, and the expression of Glutathione S-transferase Ω-1 was increased by 3.6 times, which is presumed to be caused by cellular antioxidant stress (Timmins-Schiffman et al., 2014).

It was reported that exposure of *C. elegans* to hyperosmotic conditions can also lead to up-regulation of the transcriptome of the detoxification pathway gene (glycosyltransferase, cytochrome P450, glutathione S-transferase, and ABC transporters), and their up-regulation were partially dependent on SKN-1 (Dodd et al., 2018). *skn-1* (RNAi) reduces the expression of many phase II detoxification genes, especially within the *gst* gene class (Dodd et al., 2018; Fukushige et al., 2017).There are also substances that induce up-regulation of the detoxification pathway genes independently of *skn-1*, for example, t-BOOH (tert-Butyl hydroperoxide), a stable lipid-soluble hydrogen peroxide that attacks lipids and proteins (Oliveira et al., 2009). In our study, the expression level of *skn-1* do not show significantly difference among different pH conditions (Supplementary Figure 3). Thus, the up-regulation of detoxification pathway caused by acid stress is more likely to be independent of *skn-1* regulation in *C. elegans*.

Furthermore, we found that the expression of nuclear hormone receptors genes showed similar trends with xenobiotic detoxification genes in response to acidic pH stress (Figure 5F). Nuclear hormone receptors are important transcriptional regulators involved in a variety of physiological functions, many of which are associated with diseases such as cancer, diabetes or hormone resistance syndromes. NHRs have broad substrate specificities in regulating detoxification enzymes and they are critical hubs for the metabolism of endo-and xeno-biotics (Hoffmann and Partridge, 2015). They also modulate the homeostasis of steroid hormones, other endogenous cholesterol derivatives, and lipid metabolism (Hoffmann and Partridge, 2015).

A recent study finds that *nhr-10* and *nhr-68*, both transcriptionally and functionally, are important for activating shunt gene expression in response to the excessive accumulation of propionate (Bulcha et al., 2019). In *C. elegans*, *nhr-8* has been demonstrated to regulate xenobiotic detoxification, which is necessary for anti-colchicine and chloroquine toxins (Lindblom et al., 2001). The activation of detoxification and immune response to mitochondrial toxin or pathogenic Pseudomonas aeruginosa is transcriptionally mediated by *nhr-45* (Mao et al., 2019). The NHR superfamily has conserved and typical structures, the DNA binding domain (DBD) and the ligand binding domain (LBD). Fifteen NHRs are highly conserved in *C. elegans*, Drosophila, and mammals. For example, the identity between the DBD of *C. elegans* NHR-67 and Drosophila DmTLL and human HsTLX was 79% and 70%, respectively (Maglich et al., 2001; Sluder et al., 1999). NHR-67 plays an essential role in larval development and functions as a component of a complex regulatory network that regulates the differentiation and organogenesis of vulva, thereby regulating egg laying (Fernandes and Sternberg, 2007). The identified NHR ligands are generally lipophilic including steroid hormones, glucocorticoids and exogenous drugs (Huang et al., 2010; McEwan, 2009).

In this study, the expression pattern of nhr genes is highly consistent with that of the metabolism of xenobiotics by cytochrome P450 pathway genes, indicating that the xenobiotics are likely to bind to the nuclear hormone receptors directly or indirectly through the ligands, and then activate the up-regulation of the detoxification-related genes of the xenobiotics.

Xenobitic detoxification genes are reported in other animals response to pH stress, although only one or few genes are found (Harms et al., 2014; Timmins-Schiffman et al., 2014). However, our data shows a systematic pathway genes increase in acidic stress conditions. It has been reported xenobitics detoxification genes are involved in osmotic stress (Dodd et al., 2018), heat-shock stress (Joshi et al., 2016), and hypoxia stress environment (Leiser et al., 2015). According to our data, metabolism of xenobiotics by cytochrome P450 pathway might be a universal signaling pathway to deal with stress environments. Furthermore, to manipulate xenobiotics detoxification genes through gene overexpression or CRISPR genome editing, might provide a solution to agriculture and ecosystem restoration in the global climate change contexts.

### 4.3 Down regulation of heat shock protein genes

We observed that the expression of heat shock proteins significantly down-regulated in response to acidic environments. *F44E5.4*, *hsp-70*, and *hsp-110* all encode HSP70 heat shock protein family members. As a molecular chaperone, HSP70 promotes protein folding, membrane translocation, degradation of misfolded proteins and protection of cells from stress-induced damage (Liu and Chen, 2013; Goncalves et al., 2017). HSP-110 is an ortholog of the human HSPA4. Loss of *hsp-110* activity via large-scale RNAi screens leads to a variety of defects including embryonic and larval lethality, slow growth rates, and locomotory and morphological defect (www.wormbase.org). In addition, Rampelt et al. found that knock-down of *Caenorhabditis elegans hsp110* disrupted the dissolution of heat-induced protein aggregates and severely shortened the lifespan after heat shock (Rampelt et al., 2012).

*hsp-16.1*, *hsp-16.11*, *hsp-16.2*, *hsp-16.41*, *hsp-16.48*, *hsp-16.49* all encode 16-kD heat shock protein that are members of the HSP16/HSP20 family of heat shock proteins. They are likely to function as passive ligands temporarily preventing unfolded proteins from aggregating (www.wormbase.org). HSP-16.1 and HSP-16.41 are required for an acquired tolerance to heat stroke. HSP-16.1 is localized to the Golgi and functions with PMR-1/PMR1 Ca2^+^ and Mn2^+^ transporting ATPase and NUCB-1 to maintain calcium homeostasis under heat stroke (www.wormbase.org; Kourtis et al., 2012).

Therefore, stable expression of heat shock proteins might be necessary to maintain healthy physiological conditions. In our study, the expression of heat shock protein decreases with declined pH, which is consistent with the phenotype of the negative effects of pH drop on the growth and reproduction of the nematodes.

In conclusion, our results suggested that cuticle synthesis and integrity, together with the reprogrammed xenobiotics metabolism by cytochrome P450 pathway were two major stress-responsive pathways for *C. elegans* in acidic stress environments. On one hand, when the pH drops from 6.33 to 4.33, the worms resist the damage caused by the acid culture plate by enhancing the synthesis and secretion of epidermal collagen. On the other hand, when the pH continues to decrease from 4.33, the metabolism of xenobiotics by cytochrome P450 pathway genes play a major role in protecting the nematodes from the toxic substances produced by the more acidic environment. At the same time, cuticle synthesis slows down possibly because of insufficient protective ability. To the best of our knowledge, this study is the first to systematically discover that there are two strategies which *C. elegans* respond to acidic pH stress environment: cuticle structure reorganization and xenobiotics metabolism reprogramming. Furthermore, the mechanisms found in nematode might be also applied to other invertebrate and vertebrate animals to survive in the changing pH stress environments. Thus, our data might lay the foundation to identify the key gene(s) responding and adaptation to acidic pH stress in further studies, and might also provide new solutions to improve assessment and monitoring of ecological restoration outcomes, or generate novel genotypes via genome editing for restoring in challenging environments especially in the context of acidic stress through global climate change.

## 6 Author Contributions

YC and LZ conceived and designed the experiments. YC carried out most of the experiments and analyzed the data. HY, PZ performed RT-qPCR validation. YX and XC are involved in sampling and data analysis. YC wrote the manuscript and LZ edited. LZ supervised the project. All authors read and approved the final manuscript.

## 8 Acknowledgments

We are grateful to all members of the L.Z. laboratory for their technical advices and helpful discussions.

## 9 Supplementary Material

**Supplementary FIGURE 1.**
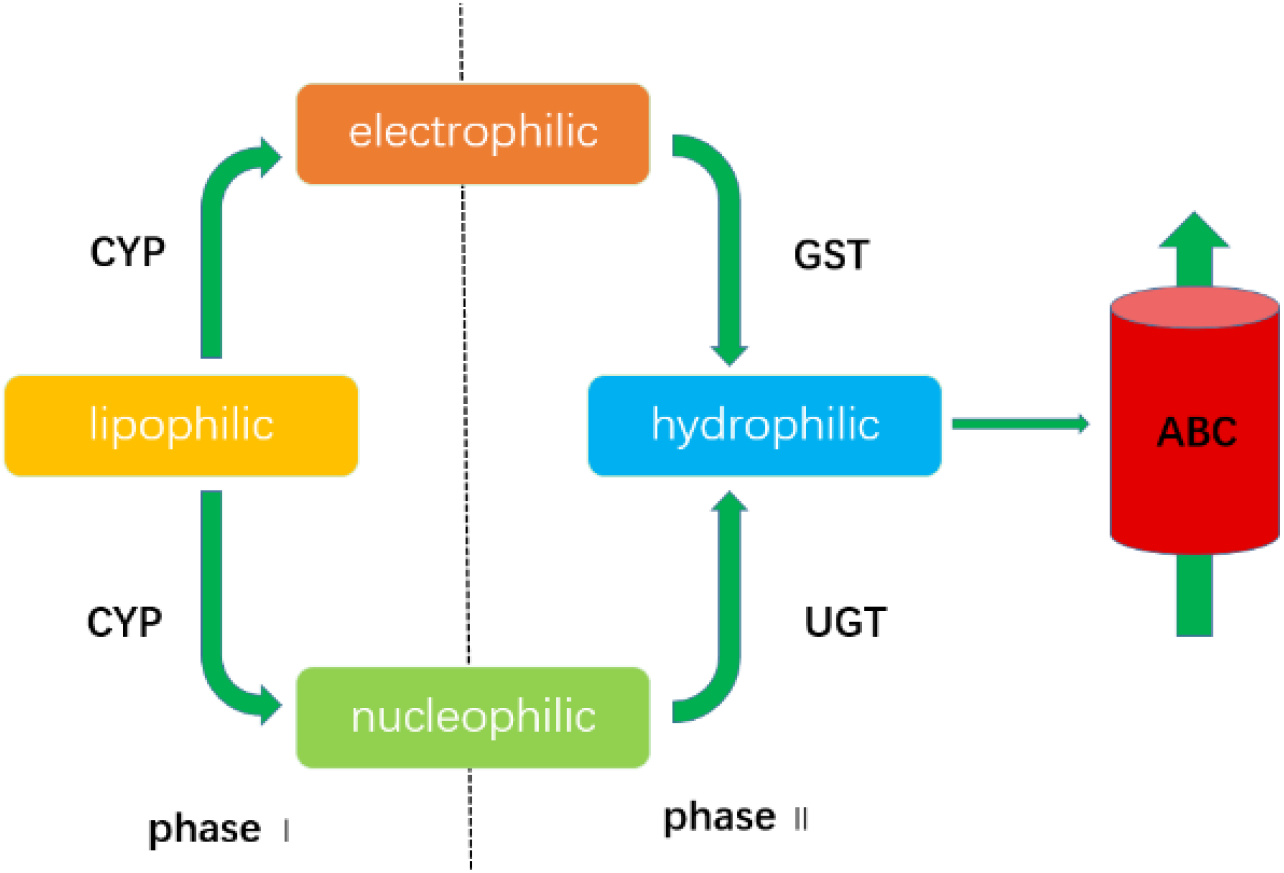
Xenobiotics metabolism by cytochrome P450 pathway. In Phase 1, lipophilic xenobiotics are metabolized to electrophilic or nucleophilic substances by phase I enzymes such as Cytochrome P450s (CYPs). In phase II, the metabolites produced by the first-stage reaction are combined with polar ligands such as glucuronic acid, glutathione etc. under the catalysis of transferases (GST, UGT, etc.). Finally, these trait-changed compounds are transported extracellularly by the ABC transporter (Barnes, 1960; Croom, 2012; Dieterich et al., 2008)

**Supplementary FIGURE 2.**
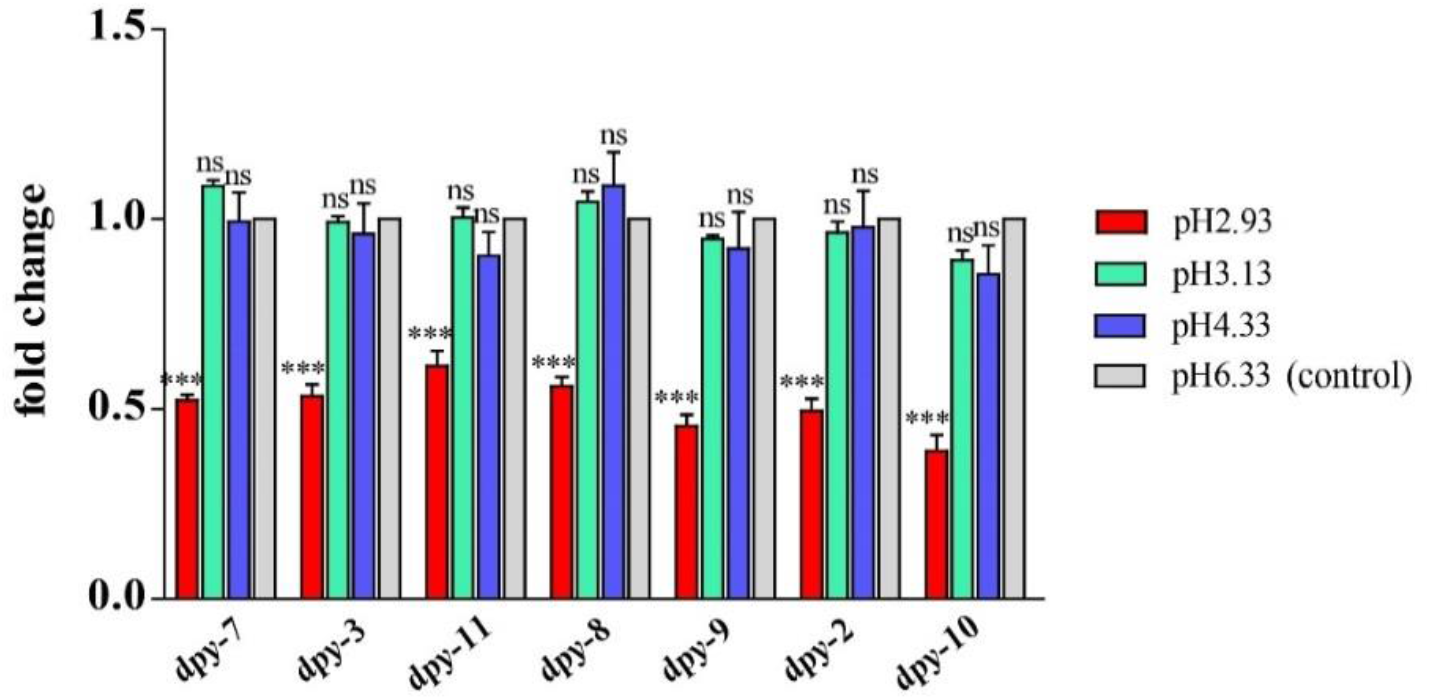
Transcript level of *dpy* genes. Fold changes indicates the ratio of the treatment group (pH 2.93, pH 3.13, pH 4.33) to the control group (pH 6.33). The error bars represent standard error of the mean of three biological replicates per condition. \**P* <0.05, \*\**P* <0.01, \*\*\**P* <0.001

**Supplementary FIGURE 3.**
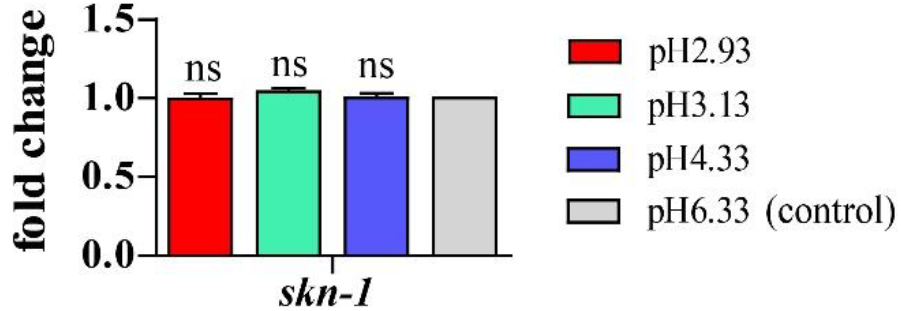
Transcript level of *skn-1* gene. Fold changes indicates the ratio of the treatment group (pH 2.93, pH 3.13, pH 4.33) to the control group (pH 6.33). The error bars represent standard error of the mean of three biological replicates per condition. \**P* <0.05, \*\**P* <0.01, \*\*\**P* <0.001

Supplementary_File_1

Supplementary_File_2

Supplementary_File_3

Supplementary_File_4

Supplementary_File_5

